# Value shapes the structure of schematic representations in the medial prefrontal cortex

**DOI:** 10.1101/2020.08.21.260950

**Authors:** Philipp C. Paulus, Ian Charest, Roland G. Benoit

**Affiliations:** Max Planck Research Group: Adaptive Memory, Max Planck Institute for Human Cognitive and Brain Sciences, Leipzig, 04103, Germany; International Max Planck Research School NeuroCom, Leipzig, 04103, Germany; School of Psychology, University of Birmingham, Birmingham, B15 2TT, UK

**Keywords:** Schema, valuation, medial prefrontal cortex, long term memory, episodic simulation, representational similarity analysis, fMRI

## Abstract

Adaptive cognition is fostered by knowledge about the structure and value of our environment. Here, we hypothesize that these two kinds of information are inherently intertwined as value-weighted schemas in the medial prefrontal cortex (mPFC). Schemas (e.g., of a social network) emerge by extracting commonalities across experiences and can be understood as graphs comprising nodes (e.g., people) and edges (e.g., their relationships). We sampled information about unique real-life environments (i.e., about personally familiar people and places) and probed the neural representations of their schemas with fMRI. Using representational similarity analysis, we show that the mPFC encodes indeed both, the nodes and edges of the schemas. Critically, as hypothesized, the strength of the edges is not only determined by experience and centrality of a node but also by value. We thus account for the involvement of the mPFC in disparate functions and suggest that valuation emerges naturally from encoded memory representations.

## Introduction

Our rich knowledge of the past allows us to readily make sense of the present. It also facilitates adaptive planning for the future, for example by supporting simulations of prospective events (1–5). Critically, these capacities are not exclusively dependent on individual memories of unique past experiences. Instead, they are also based on generalized knowledge about our environment that is derived from multiple experiences (e.g., knowledge about relationships between familiar people)(3, 6).

A type of such generalized knowledge structures are memory schemas (7–9). These representations of our environment can be understood as graphs comprising information about nodes (e.g., individual people) and their edges (e.g., their relationships) (8, 10–12). Schemas are formed by extracting commonalities across related events (7, 8, 13). They thereby reduce the complexity of our experience into simplified models of the world (e.g., about the people we know or about the locations we frequently visit) (8, 14). Such models, in turn, foster planning and facilitate adaptive decisions (15–17).

However, beyond a representation of the environment’s structure, adaptive cognition also requires a representation of what’s valuable within that environment (18). Here, we test the hypothesis that these two kinds of information are inherently intertwined in the rostral and ventral medial prefrontal cortex (mPFC) (19–22). As detailed below, this proposal accounts for the involvement of this region in two seemingly disparate functions: representing *memory schemas* and *value*.

Evidence from humans (23, 24) and rodents(20, 25) indicates a critical role for the mPFC in mediating *memory schemas* (26, 27). Activity patterns in this region have been shown to code for individual nodes of the environment, such as for familiar people (19, 28) and places(19, 29). However, it remains unclear whether the mPFC encodes representations of the nodes in isolation or whether these representations also entail information about their edges (i.e., their relationships to other nodes).

A largely independent line of research has associated the mPFC with the representation of affect and *value* (21, 30– 32). Activity in this region tracks the value of objects, people, or places that we currently perceive or imagine (19, 30, 33– 35). Moreover, in humans, focal lesions disrupt value judgements (36, 37). The mPFC has thus been argued to represent value in a common currency that allows for flexible decision making in a wide range of contexts (30, 31, 38). Notably, evidence from human neuroimaging (1, 19, 33, 35, 39) and rodent single cell-recordings (20, 22) has shown that representations of memories and of value are supported by *overlapping* parts of the mPFC. We thus reconcile the common attribution of these functions by hypothesizing that the schemas encoded by this region are shaped by value.

Specifically, we propose that the mPFC encodes representations of individual nodes (e.g., individual familiar people) and that the representations also entail information about their edges (e.g., the overall associations between the people). Critically, we suggest that nodes that are more important for a person exhibit stronger edges.

We hypothesize that the importance of a given node is jointly determined by three features: Given that schemas build up with experience (8), we first expect that more familiar nodes should be more prominently embedded in the overall graph (33). Secondly, for the same reason, we expect stronger embedding of nodes that are more central to the respective environment (11). Finally, given the role of the mPFC in affect and valuation (21, 30), we propose the edges are also weighted by the nodes’ value (19, 20, 40). The encoded schemas would thus emphasize connections of behaviorally relevant elements of the environment, reminiscent of the hippocampal weighting of rewarded locations (41, 42).

Here, we test this hypothesis by probing the neural representations of two distinct and individually unique schemas: about people’s social networks and about places from their everyday environment. This allows us to examine whether the suggested coding principles generalize across these individual schemas. Participants provided names of people and places they personally know and arranged these names in circular arenas according to their associations (43). This allowed us to quantify the centrality of each exemplar (e.g., a person) to its respective schema (e.g., the social network). Participants also indicated their familiarity with each person and place (as an index of experience) and their liking of each of these exemplars (as an index of affective value). In a subsequent session, we measured their brain activity using functional magnetic resonance imaging (fMRI) while they imagined interacting with each person and being at each place. We took the ensuing activity patterns to assess the neural representations of the individual nodes and their edges using representational similarity analysis (RSA) (44).

First, we hypothesized that the mPFC encodes unique representations of the nodes that get reinstated during mental simulation (19, 28, 29). We thus predicted similar activity patterns to emerge in the mPFC whenever participants imagine the same person or place. Second, we hypothesized that the structure of neural activity patterns *across* nodes reflects the structure of their edges. That is, we reasoned that pairs of nodes that are more strongly connected (i.e., that exhibit stronger edges) are encoded by more overlapping neuronal populations (1, 45–47). This, in turn, should be reflected in overall greater neural similarity for nodes with particularly strong edges. As a consequence, if more important nodes have stronger edges, they should also exhibit overall greater neural similarity.

In addition, we further gauge the regional specificity of such value-weighted schemas to the mPFC. Therefore, we also examine the posterior cingulate cortex and the hippocampus, two regions that have similarly been associated with memory (48–51) and valuation (30, 34, 52).

## Results

### The medial prefrontal cortex codes for the nodes of participants’ real-life schemas

We first examined the hypothesis that the medial prefrontal cortex encodes representations of personally familiar people and places, i.e., the nodes of the respective schemas. Whenever we simulate an event involving a particular node, its representation should get reinstated in the mPFC. We thus took the ensuing fMRI activity patterns as proxies of their respective neural representations (19, 53) and examined their replicability using an RSA searchlight approach (radius = 8 mm, 4 voxels) (44).

In regions that encode the nodes of the schema, we predicted overall greater pattern similarity for simulations featuring the same node (same-item similarity) than for simulations featuring different nodes (different-item similarity) (54). Note that the different-item measure was only based on the similarity of activity patterns for nodes of the same kind (i.e., either people *or* places). This ensured that the results are not influenced by potential categorical differences in the representation of people versus places (Figure 1A and 1B) (19, 53).

**Figure 1.**
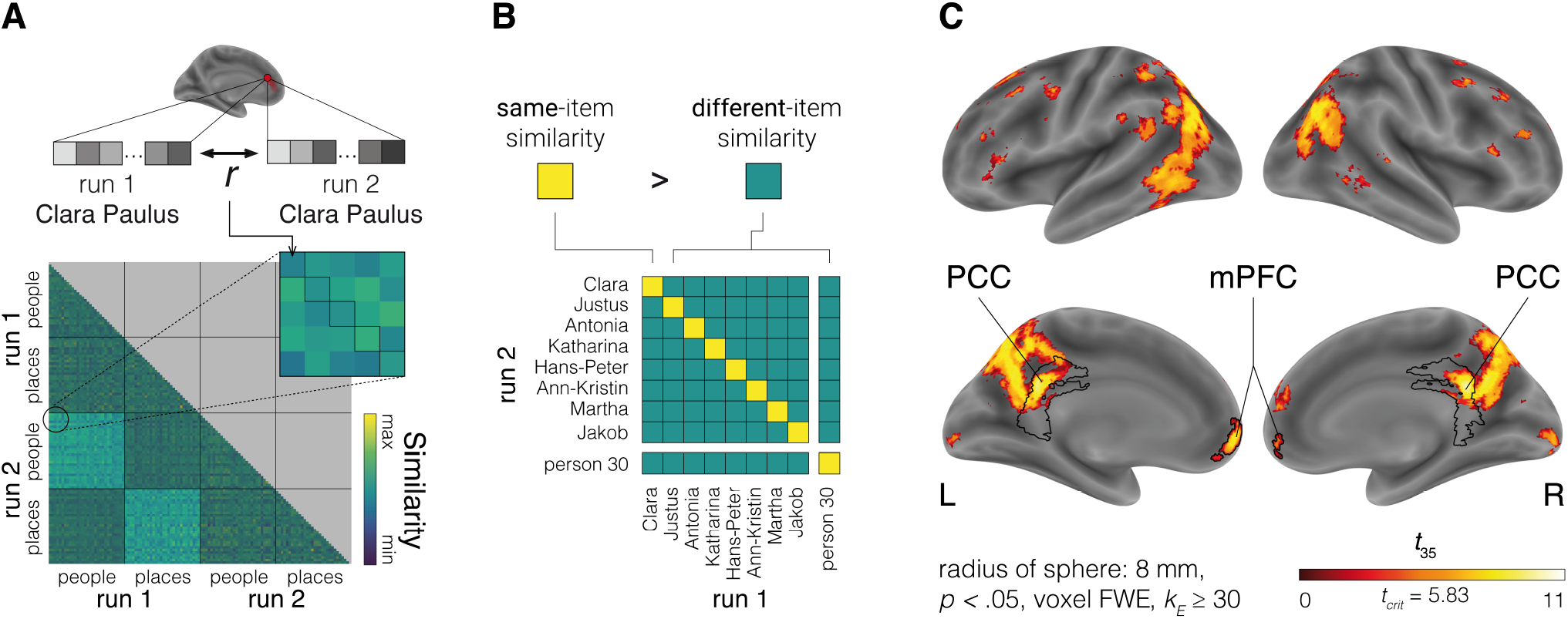
Representations in the mPFC and PCC code for the nodes of participants’ real-life schemas. (A) We examined whether the mPFC encodes representations of the nodes by testing for the replicability of activity patterns for the same people and places across the two functional runs. Each row and column of the representational similarity matrix corresponds to a single simulation trial. (B) Regions coding for the nodes should show more similar activity patterns for the repeated simulations of the same person or place (*same-item similarity*) than for simulations entailing different nodes of the same category (*different-item similarity*). (C) The searchlight analysis identified regions coding for the nodes of real-life schemas. These entailed the mPFC and PCC. mPFC = medial prefrontal cortex, PCC = posterior cingulate cortex.

Corroborating our previous finding (19), we obtained this effect in the mPFC. This region thus yielded replicable activity patterns that were specific to individual exemplars (Figure 1C, Table S1). Moreover, we also observed evidence for such replicable pattern reinstatement in a number of other brain regions that are typically engaged during the recollection of past memories and the simulation of prospective events (50, 55, 56). These regions included the posterior cingulate cortex (PCC), the precuneus and parts of the lateral parietal and temporal cortices. Notably, there was no evidence for pattern reinstatement in the hippocampus.

The data thus support our hypothesis that the mPFC encodes unique representations of individual nodes. In the following, we further examine the edges *between* nodes in the mPFC and PCC regions of interest (ROI) identified by this analysis. We also test for these effects in the hippocampus, even though this region showed no evidence of node coding.

Note that the subsequent analyses of the edges are based on different parts of the neural representational similarity matrix (RSM) than the ones used to determine node coding. Further, they are based on comparisons of model RSMs that are also independent of the node coding model. In the supplement, we provide complementary and consistent results based on anatomically defined masks (see Tables S2, S3, S4; Figure S1 and S2).

### Centrality, experience, and affective value load on a common principal component that quantifies importance

We had hypothesized that a node’s centrality, experience, and also its affective value contribute to its importance. These three features may thus share a common latent factor. First, to assess *centrality*, participants positioned the names of the people and places in circular arenas (Figure 2A). They were instructed to arrange nodes close together if they associate them strongly with each other and far apart if they do not (43). We calculated the centrality of each node as the sum of its inverse distances to all other nodes. Participants then arranged the people and places on continuous scales providing estimates of their familiarity with each node (as an index of *experience*) and of their liking (as an index of *affective value*). All three features were assessed separately for people and places.

**Figure 2.**
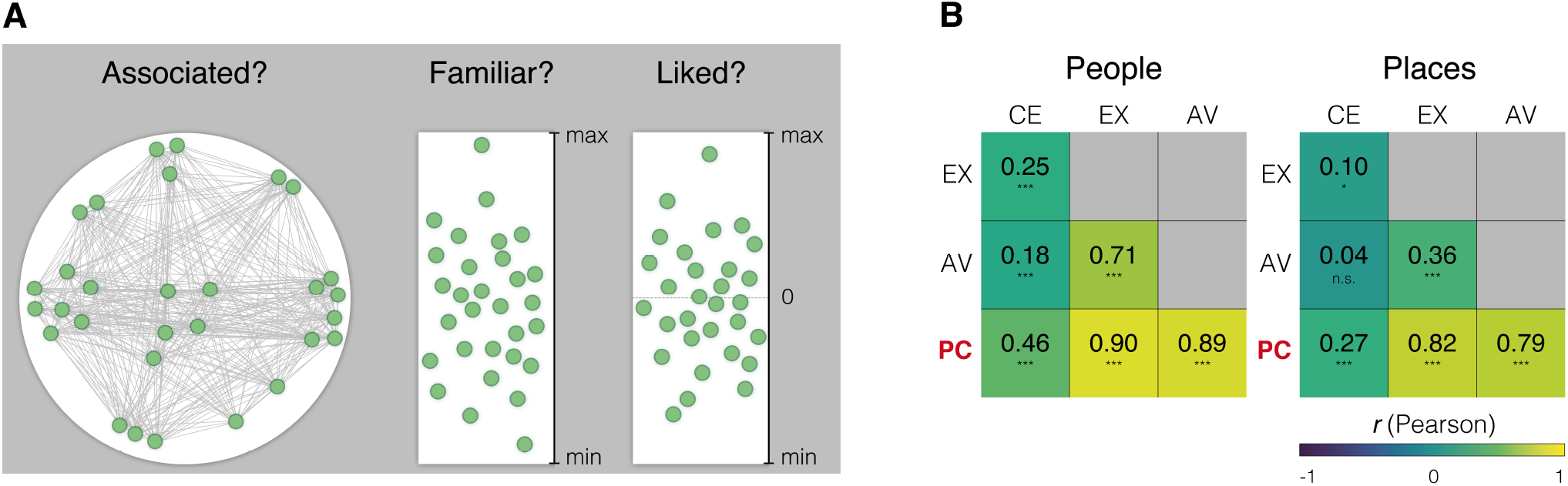
Centrality, experience, and affective value load on a common principal component that quantifies importance. (A) Participants arranged the familiar people and places on circular arenas according to their associations, thus providing a measure of centrality. Participants also provided measures of experience and affective value by indicating their familiarity with the people and places as well as their liking. (B) Centrality, experience, and affective value load on a common principal component as indicated by significant positive correlations (* - *p* < .05, *** - *p* < .001; one-tailed; *df* = 35). This component thus summarizes the importance of the nodes (i.e., the people and places) to the schema. CE = centrality, EX = experience, AV = affective value, PC = principal component.

To test whether centrality, experience, and affective value load on a common latent factor, we *z*-scored each vector of values separately for each category (people, places) and within each participant. This approach prevents between-participant variance from influencing the factor solution. We then performed principal component analyses, separately for the people and places. The respective first principal component explained, across all participants, 61% of variance for people and 46% for places.

Critically, as predicted, both of these principal components exhibited significant positive correlations not only with experience and centrality but also with affective value (Figure 2B). We thus take them to quantify the importance of each individual node to its respective schema. In the next step, we used the individual importance values of the respective principal component to predict the structure of the schemas’ edges.

### The medial prefrontal cortex encodes the edges of value-weighted schemas

We had hypothesized that more important nodes – as indicated by the principal component – exhibit stronger edges. We had further reasoned that the strength of edges is reflected in the neural similarity of the connected nodes. Specifically, we assumed that more strongly connected nodes are also encoded by more overlapping neuronal populations (1, 20, 57). As a consequence, we had predicted that more important nodes should exhibit overall greater neural similarity.

We tested this prediction by constructing models of the expected structure of representations in the mPFC. The models were based on the importance values derived from the respective principal component. Specifically, we predicted the similarity between any two nodes by the product of their respective principal component scores (i.e., importance values). Thus, we expected more important nodes to yield overall greater pattern similarity (Figure 3B).

**Figure 3.**
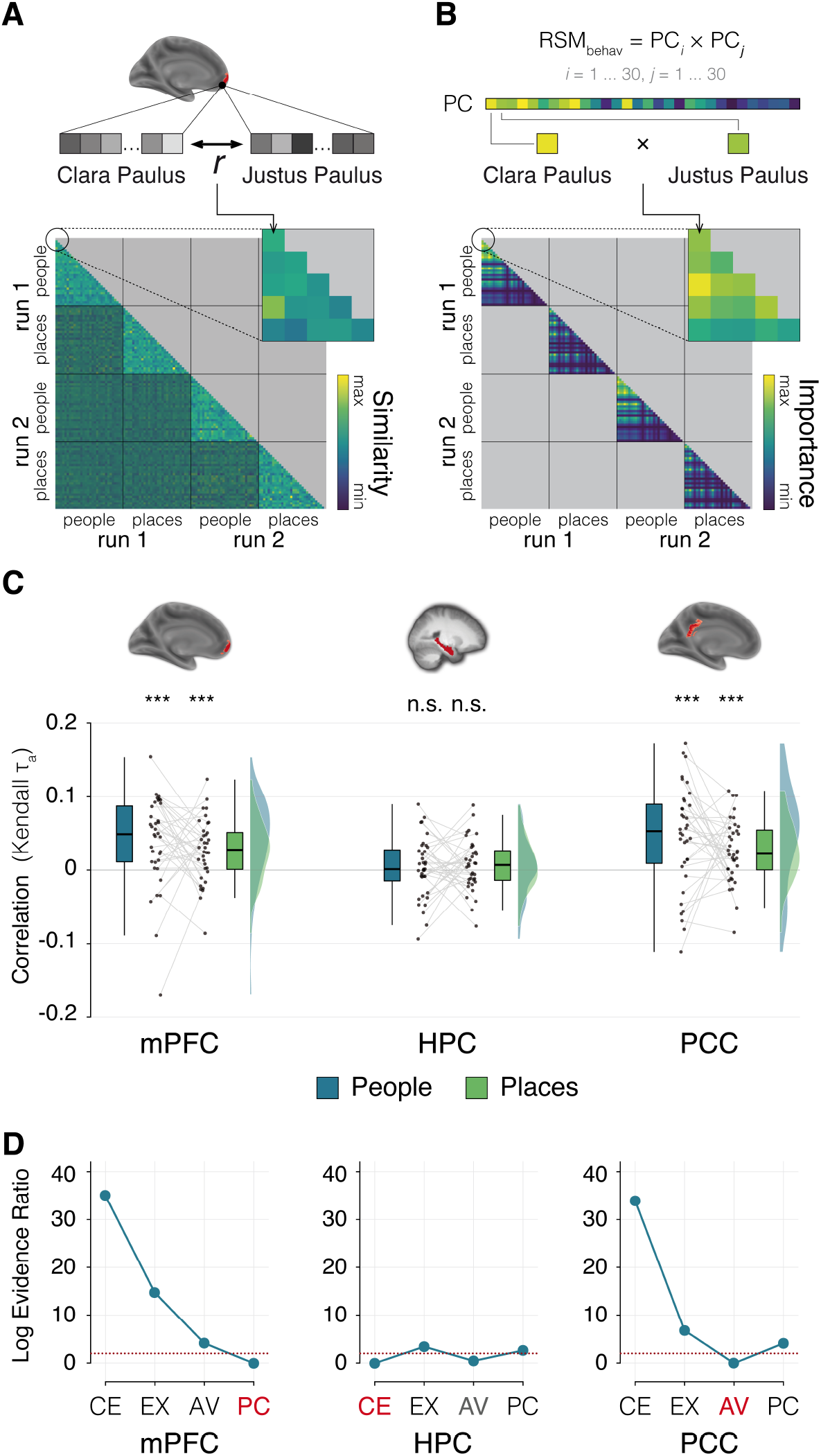
Only the structure of representations in the mPFC is best accounted for by the principal component model that reflects importance. (A) Construction of the neural RSM. Each row and column of the matrix corresponds to a single simulation trial. In this analysis, we examine the similarity of activity patterns elicited by simulations of different people or places. (B) Construction of the model predictions. We predicted more similar representations for people and places with overall higher principal component scores, given that these more important exemplars should be more strongly embedded in their overall schema. To this end, we computed the combined importance of any two people or places from the product of their principal component scores. (C) Correlation of neural RSM and model prediction. Asterisks denote significant positive correlations as tested in a *t* -test on the *Fisher-z* transformed correlation coefficients (*** - *p* < .001; one-tailed; *df* = 35). Box-plots: center line, median; box limits, first and third quartile; whiskers, 1.5x interquartile range. (D) Comparisons of linear mixed models further support the hypothesis: only the structure of representations in the mPFC is best explained by the principal component. The figure displays Log Evidence Ratios (LER). Smaller values indicate better fit. By definition, the best model assumes a value of zero. The dotted red line demarks a relative LER difference of two, regarded as decisive. mPFC = medial prefrontal cortex, HPC = hippocampus, PCC = posterior cingulate cortex, CE = centrality, EX = experience, AV = affective value, PC = principal component.

We then determined the neural similarity structure in the mPFC, PCC, and in the hippocampus (Figure 3A). We constrained the broader cluster containing the PCC to the parts covering this region using an anatomical mask (58). We used an anatomical mask from the same atlas to examine the representational structure for the hippocampus. All analyses were conducted in subject space.

Finally, we tested for the correspondence between our prediction and the actual structure of neural representations by computing the correlation of the respective parts of the lower triangular vectors of both matrices (Figure 3C). This was done separately for people and places to examine whether the effect is present for either category. Using Kendall’s *τ*_*a*_ as a conservative estimate (59), we indeed observed significant correlations in the mPFC for both people (*mean τ*_*a*_ = 0.039, tested with a Wilcoxon test, *W* = 562, *p* < .001, *d* = 0.63, one-tailed, due to a deviation from normality indicated by a Shapiro-Wilk test, *W* = 0.92, *p* = 0.01) and places (*mean τ*_*a*_= 0.026, *t*(35) = 3.72, *p* < .001, *d* = 0.62, one-tailed) - with no significant differences between the two (*t*(35) = 1.04, *p* =.307, *d* = 0.17, two-tailed).

Similarly, the correlations were also significant in the PCC for people (*mean τ*_*a*_ = 0.046, *t*(35) = 3.91, *p* < .001, *d* = 0.65, one-tailed) and places (*mean τ*_*a*_ = 0.026, *t*(35) = 3.61, *p* < .001, *d* = 0.60, one-tailed), again with no significant differences between the two (*t*(35) = 1.42, *p* = .165, *d* = 0.24, two-tailed). However, the same analyses of the hippocampal data did not yield evidence for a match between the predicted and actual structure of representations (people: *mean τ*_*a*_ = 0.003, *t*(35) = 0.41, *p* = .341, *d* = 0.07, one-tailed; places: (*mean τ*_*a*_ = 0.007, *t*(35) = 1.15, *p* = .13, *d* = 0.19, one-tailed). We obtained qualitatively identical results in analyses based on purely anatomically defined ROIs (see Table S2 and S3; Figure S1). The results are also in accordance with a whole-brain searchlight analysis (radius = 8 mm, 4 voxels) (see Table S5). We thus show that representations in the mPFC generally align with the predicted structure of value-weighted schematic representations. However, it remains to be determined whether importance is indeed the best model to account for the structure of representations in any of our ROIs.

### The importance model accounts best for the structure of mPFC representations only

Does the structure of representations predicted from the principal component account best for the data or would any of the individual contributing features provide at least a comparable or even a better fit? If the mPFC does encode value-weighted schemas, we would expect the model based on the conglomerate index of importance to outperform models only based on centrality, experience, or affective value. Furthermore, we would expect some degree of regional specificity, i.e., that only representations in the mPFC are best accounted for by importance.

We formally tested these predictions by setting up alternative models that were solely based on either centrality, experience, or affective value. We then compared these models with the importance model that was based on the principal component. In brief, we set up linear mixed effects models to account for the structure of representations as a function of each of these individual features. In each of these models, we included a factor of category (people, places) and the maximum possible random effects structure that would converge across all models and regions of interest. We thus accounted for between participant variance by including a random intercept per participant and run, as well as random slopes for our fixed effects of category and the respective predictor (e.g., the principal component scores). We then performed model comparisons within each ROI to determine the model that best fits the neural similarity structure. The comparisons were based on Log Evidence Ratios (*LER*) derived from Akaike’s Information Criterion (60). We regarded *LER* differences greater than two as decisive evidence for the better model (61).

Consistent with our hypothesis, in the mPFC, the principal component model accounted best for the data. It performed decisively better than affective value (*LER* = 4.19), experience (*LER* = 14.8) and centrality (*LER* = 35.16) (Figure 3D). The model parameters of this winning model entailed a significant main effect of category, reflecting overall higher neural pattern similarity for people than places (*β*_*Category_place*_ = -0.026, *SE* = 0.008, *χ*^2^ = 11.61, *p* < .001). Critically, they also included a significant positive parameter estimate for the principal component, indicating overall greater neural pattern similarity for nodes of greater importance (*β*_*PrincipalComponent*_ = 0.048, *SE* = 0.012, *χ*^2^ = 17.12, *p* < .001). Moreover, the main effect of the principal component did not interact with category (*β*_*Category_place*:*P rincipalComponent*_ = -0.005, *SE* = 0.008, *χ*^2^ = 0.36, *p* = .546).

By contrast, in both control regions, other models were better suited to account for the structure of representations. For the hippocampus, the model comparison yielded the best fit for centrality, though there was only a minimal advantage for this model over affective value (*LER* = 0.49). Notably, both models performed decisively better than the ones based on either the principal component (*LER* = 2.68) or experience (*LER* = 3.46). However, of the model parameters, only the main effect of category was significant, indicating overall higher pattern similarity for places than for people (*β*_*Category*_*place*_= 0.02, *SE* = 0.004, *χ*^2^ = 31.49, *p* < .001). There was neither a main effect of centrality (*β*_*Centrality*_ = -0.007, *SE* = 0.004, *χ*^2^ = 0.9, *p* = .342) nor an interaction of category with centrality (*β*_*Category_place*:*Centrality*_ = 0.007, *SE* = 0.004, *χ*^2^ = 3.04, *p* = .081). The same pattern (i.e., only a main effect of category) also emerged for the model based on affective value (see Table S4).

For the PCC, the model based on affective value performed decisively better than any other model: principal component (*LER* = 4.14), experience (*LER* = 6.82), and centrality (*LER* = 33.99) (Figure 3D). The main effect of affective value was significant, indicating overall greater neural pattern similarity for nodes of higher affective value (*β*_*AffectiveV alue*_ = 0.026, *SE* = 0.007, *χ*^2^ = 10.71, *p* = .001). There was no main effect of category (*β*_*Category_place*_ = 0.014, *SE* = 0.009, *χ*^2^ = 1.51, *p* = .219), but an interaction of affective value with category, reflecting a stronger effect of affective value for people than places (*β*_*Category_place*:*Af fectiveV alue*_ = -0.01, *SE* = 0.005, *χ*^2^ = 3.92, *p* = .048) (see Table S4 for all model parameters).

In summary, the model based on the principal component was the clear winner in the mPFC, whereas it was outperformed by alternative models in the control regions. This pattern thus suggests some regional specificity of value-weighted schemas. Note that we obtained consistent results when examining the structure of representations in the purely anatomically defined ROIs (see Figure S2; Table S4).

Finally, we sought to ensure that the winning models in each ROI perform better null models based on noise. To this end, for each familiar person and place, we randomly sampled a value from a standard normal distribution. We then used these values to construct a noise model by performing the identical processing steps as for the other predictors.

Moreover, this allowed us to derived a second noise model by first sorting the vector of noise values in descending order prior to constructing the model. This model mirrors the order in which participants tend to list people and places, i.e., by starting with people and places they like and know better. As a consequence, nodes that are listed earlier tend to also have higher values on the principal component. By sorting the noise vectors in descending order, we imposed a similar dependence between the noise values and their serial positions (Figure 4A).

**Figure 4.**
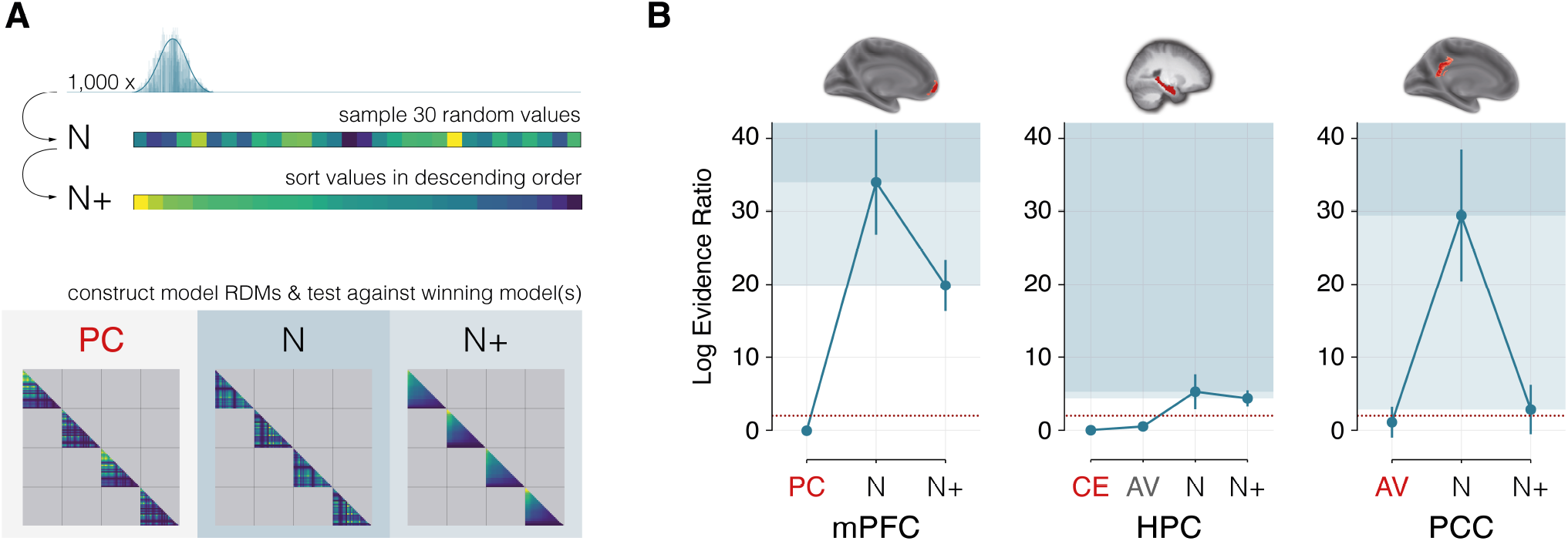
The importance model outperforms noise models in the Mpfc. (A) Construction of the noise models. On 1,000 iterations, we sampled 30 values for each category type from a standard normal distribution to create a vector of noise values. We then used these vectors to construct a random noise model (N) by performing the same processing steps as for the other models. This approach also allowed us to create a sorted noise model (N+). Here, we first sorted the vector of noise values in descending order. This model mirrors the order in which participants tend to list people and places, i.e., by starting with ones that are more familiar and liked. (B) Comparisons with noise null models. Points depict the mean model performance across comparisons with 1,000 random noise models (N) and sorted noise models (N+), whiskers indicate the standard deviation. Smaller values indicate better fit. The dotted red line demarks a relative LER difference of two, regarded as decisive. mPFC = medial prefrontal cortex, HPC = hippocampus, PCC = posterior cingulate cortex, CE = centrality, EX = experience, AV = affective value, PC = principal component, N = random noise, N+ = sorted noise.

Separately for each ROI, we then compared model performance of random noise and sorted noise against the winning model. Because there was no clear single winning model for the hippocampus, we included the models based on centrality and on affective value (*LER* < 2) in this comparison. We repeated this process 1,000 times to obtain an estimate of the expected performance of the noise models and the winning model. As expected, in the mPFC, the principal component remained the best model (*mean LER* = 0, *SD* = 0.04), performing decisively better than sorted noise (*mean LER* = 15.56, *SD* = 2.51) and random noise (*mean LER* = 25.81, *SD* = 6.03).

For the hippocampus, the initial model comparisons had provided only minimal evidence for centrality over affective value. We therefore compared both of these models with the noise models. Again, there was minimal evidence for superiority of centrality (*mean LER* = 0.09, *SD* = 0.53) over affective value (*mean LER* = 0.58, *SD* = 0.53). Both models performed decisively better than sorted noise (*mean LER* = 4.4, *SD* = 1.09) and random noise (*mean LER* = 5.32, *SD* = 2.38). The model comparisons in the PCC revealed strong, though not decisive, evidence for a superiority of affective value (*mean LER* = 1.15, *SD* = 2.1) over sorted noise (*mean LER* = 2.89, *SD* = 3.37). However, both did perform decisively better than random noise (*mean LER* = 29.46, *SD* = 9.01) (see Figure 4B).

The results thus provide further evidence for the hypothesis that the mPFC encodes both the nodes and the edges of value-weighted schematic representations. The model comparison moreover supports this account with some regional specificity.

## Discussion

Human adaptive cognition is fostered by representations of the structure of our environment (7, 15, 17). Such structured representations act as templates that allow us to facilitate recollections of the past, to make sense of the present, and to flexibly anticipate the future (6, 8, 13, 33, 62). Structured representations have been described in the mPFC for various domains, ranging from spatial and conceptual to abstract state spaces (63–67).

Our results support the hypothesis that the mPFC supports a specific form of such structured representations: value-weighted schemas of our environment. Generally, the mPFC has long been argued to mediate memory schemas (26, 27, 62), yet the exact contribution of this region has remained unclear. It has been suggested that the mPFC serves to detect congruency of incoming information with schematic knowledge that is represented in posterior areas (27). This region would thus not necessarily represent any kind of schematic knowledge by itself. Our data indicate that the contribution of the mPFC goes beyond congruency detection: It directly encodes schematic representations of the environment (11, see also 19).

These representations could act as pointer functions that guide the reinstatement of relevant distributed information (26, 27, 68, 69). This suggestion fits with broader accounts that situate the mPFC on top of a cortical hierarchy as a convergence zone (48, 70) that integrates information from diverse brain networks (33, 55).

Critically, our results support the hypothesis that schematic representations in the mPFC (e.g., of one’s social network) inherently entail the value of the encoded nodes (e.g., how much we like individual people). That is, the structure of the edges could best be accounted for by a model based on a latent factor that quantifies the importance of the nodes. As predicted, this factor was not only influenced by the nodes’ centrality (11, 40) and familiarity (19, 29, 33), but also by their affective value (19–21). We obtained this pattern across schemas for personally familiar people and places. The convergent results thus demonstrate that this coding scheme in the mPFC generalizes to different kinds of environmental representations.

The model comparison also suggested some degree of regional specificity for value-weighted schemas. The importance model was neither the best fit to the data obtained from the PCC nor from the hippocampus. Whereas even the best model did not decisively outperform a noise model in the PCC, there was some evidence that the structure of the edges in the hippocampus could best be accounted for by either centrality or affective value. These results are consistent with evidence showing that the hippocampus encodes map-like representations of relational abstract (45, 71) and social (72) knowledge and that it is involved in value learning (73, 74).

More broadly, a functional dissociation between the hippocampus and mPFC is also consistent with the suggested involvement of these regions in two complementary learning systems. Whereas the hippocampus is critical for the retention of individual episodes, the mPFC may extract commonalities across similar events and bind these into consolidated representations (13, 17, 41, 75, cf. 76). The mPFC would thus reduce the complexity of our experience into schematic summary representations.

Indeed, a recent study provided convergent evidence for such dimension reduction in this region. It demonstrated that the mPFC compresses rich perceptual input to only those features that are currently task-relevant – akin to a principal component analysis (14). While such dimension reduction entails the loss of specific details, it also affords generalizability and cognitive flexibility (8, 77). These representations can thus augment planning (6, 78, 79) and also be flexibly used for the construction and valuation of novel events (1, 33).

The emergence of schemas in the mPFC could be fostered by hippocampal replay of past events (80–82). Such replay, conveyed by monosynaptic efferent projections into the mPFC (83), can potentially provide a teaching signal that facilitates neocortical consolidation (41). Moreover, to the degree that replay is biased towards valuable information, it may lead to a stronger weighting of those experiences that are of particular importance (41, 42). However, the mPFC likewise receives direct projections from areas such as the amygdala and the striatum (84) that could also contribute to a shaping of the schematic representations by value (see also 21).

Importantly, the highlighted structure of representations in the mPFC provides a common account for the involvement of this region in both memory schemas and valuation. That is, when we think about an individual element from our environment (e.g., a known person), its representation in the mPFC is activated. This activation then spreads throughout the network of connected nodes. Critically, we suggest that there is a wider spread from nodes that are more valuable and that are thus more strongly embedded in their overarching schema. This wider spread, in turn, may manifest as greater regional univariate activity. According to this account, the valuation signal that has been attributed to the mPFC (30, 34) thus constitutes an emergent property of the structure of its encoded representations (85).

This interpretation similarly accounts for the stronger engagement of the mPFC when individuals think about themselves as compared to others (86, 87). The self can be considered a super-ordinate schema that entails abstracted representations of all our personal experiences (68). Instantiating this schema would thus presumably lead to wide spread activity, whereas thinking about specific other people would only co-activate neural representations of more restricted nodes. Moreover, the net activity would be lower for other people that we feel less connected to and that we have less experience with (33, 88–90).

To conclude, this study provides evidence that the medial pre-frontal cortex represents the structure of our environment in the form of value-weighted schemas. These schemas reflect our experience with individual nodes as well as their centrality. Critically, they also inherently encode information about their affective value. These schematic representations thus prioritize information that is critical for adaptive planning and ultimately promotes our well-being and survival.

## Methods

### RESOURCE AVAILABILITY

#### Lead contact

Further information and requests for resources should be directed to and will be fulfilled by the Lead Contact, Philipp C. Paulus.

#### Materials availability

This study did neither use nor generate new materials.

#### Data and code availability

Participants did not give consent for their MRI data to be released publicly within the General Data Protection Regulation 2016/679 of the EU. We can thus only share data with individual researchers upon reasonable request. The second level *t*-map for the node coding searchlight analysis (Figure 1C) and the functional masks of the mPFC and PCC ROI are available at neurovault: https://identifiers.org/neurovault.collection:8129

Custom code is publicly available via the Open Science Framework: https://osf.io/6h58e/?view_only=b21fa36180a845e69671f222a110bac8

### EXPERIMENTAL MODEL AND SUBJECT DETAILS

#### Participants

We recruited 39 right handed healthy unmedicated adults (23 females; *mean* age = 25.4 years, *SD* = 2.6 years) from the study database of the Max Planck Institute for Human Cognitive and Brain Sciences. All participants had normal or corrected to normal vision, provided written informed consent and received monetary compensation for their participation. The experimental protocol was approved by the local ethics committee (Ethical Committee at the Medical Faculty, Leipzig University, Leipzig, Germany; Proposal number: 310/16-ek). Three participants had to be excluded from analysis either because of a recording error (*n* = 1), or excessive movement (*n* = 2). Excessive movement was defined as absolute movement ≥3 mm within either run or a total of ≥5 episodes of movement ≥0.5 mm. We thus included 36 participants (22 females; *mean* age = 25.2 years, *SD* = 2.5 years) in the analyses.

### METHOD DETAILS

#### Task and procedures

The procedure, adapted from ref. 19, comprised two sessions. During the first session, participants provided names of personally familiar people and of such places. Participants tend to start by listing people and places that they are most familiar with and that they like the most. We therefore asked them to provide us with 90 people and 90 places and then randomly sampled 30 of each to ensure a greater variability in these variables of interest.

#### Arrangement tasks: Assessing the schema

To quantify the centrality of each node to its schema, participants arranged the names of the people and places on separate two-dimensional circular arenas using the multiple arrangements task (43) (Figure 2A). We instructed participants to position names closer to each other that they also associate more strongly. The inverse of the distance thus serves as a measure of associatedness between any two nodes. We quantified the centrality of each person and place to its schema by computing their centrality, i.e., the sum of their associatedness values.

We then assessed how much experience participants had with each person and place. The participants therefore placed the names on continuous familiarity scales ranging from “not at all familiar” to “very much familiar”. Finally, participants provided a measure of affective value for each person and place by arranging their names on continuous liking scales ranging from “not at all liked” to “very much liked” (Figure 2A). All arrangements were done separately for people and places.

#### Simulation task: Assessing neural representations

The participants returned for a separate session (*median* delay: 1 day; *range*: 1–4 days) to complete the episodic simulation task in the fMRI scanner. Each trial of the simulation task began with a fixation period of 0.5 s followed by the name of a person or a place for 7.5 s. During this time, participants imagined interacting with the person in a typical manner or being at the place engaging in a location specific activity. Participants were instructed to imagine the episode as vividly as possible, so that they have a clear mental picture of the respective person or place. Participants then rated the vividness of their imagination on a five-point scale within a maximum of 3 s. Trials for which participants failed to press a button within that time period were later removed from analysis. If there was time left from the response window, it was added to the subsequent inter trial interval. This lasted for at least 3 s plus an additional jittered period (0 to 8 s in 2 s intervals). The screen during the inter trial interval was blank. Each person and place was presented once in each of the two functional runs that followed different random orders. Before entering the scanner, participants practiced the simulation task with people and places that they had previously provided but that did not feature in the simulation task proper.

After the simulation task, participants were presented with people-places and faces-places localizers. Outside the scanner, they provided further information regarding the associations and identities of the individual people and places, including their addresses and locations. They also completed a number of standard questionnaires. These data were not analyzed for the current study.

### QUANTIFICATION AND STATISTICAL ANALYSIS

#### fMRI data acquisition

Participants were scanned with a 3 Tesla Siemens Magnetom PRISMA MRI scanner with a 32-channel head coil. We acquired anatomical images with a T1-weighted magnetization-prepared rapid gradient-echo sequence (MPRAGE, 256 sagittal slices, TR = 2,300 ms, TE = 2.98 ms, flip angle = 9°, 1 x 1 x 1 mm^3^ voxels, FoV = 240 mm, GRAPPA factor = 2). For each of the two functional runs of the simulation task, we acquired 469 volumes of blood-oxygen-level-dependent (BOLD) data with a T2*-weighted echo-planar imaging (EPI) pulse sequence (91, 92). This sequence employed multiband RF pulses with the following parameters: 72 interleaved axial-oblique slices (angled 15° towards coronal from AC-PC), TR = 2,000 ms, TE = 25 ms, flip angle = 90°, 2 x 2 x 2 mm^3^ voxels, 6/8 partial Fourier, FoV = 192 mm, MF = 3). The first five volumes of each run were discarded to allow for T1 equilibration effects.

#### Pre-processing

Data were analyzed using SPM12 (93)(www.fil.ion.ucl.ac.uk/spm) in Matlab (version 9.3). The functional images were corrected for slice acquisition times, realigned, corrected for field distortions, and co-registered with the anatomical scan. Correction for field distortions was achieved using FSL topup (94, 95) as implemented in FSL (https://fsl.fmrib.ox.ac.uk/).

#### General linear model

We then decomposed the variance in the BOLD time-series using a general linear model (GLM) in SPM12 (93). Each model included six regressors representing residual movement artifacts, plus regressors modeling the intercepts of block and session. The additional regressors in the GLM coded for the effects of interest.

Specifically, we modeled each trial as a separate condition yielding a total of 120 regressors – one for each of the two simulations of the 30 people and 30 places. The trial regressors were convolved with the canonical hemodynamic response function. A 1/128-Hz high-pass filter was applied to the data and the model. We computed *t*-maps for the estimated parameters of interest (i.e., for each simulation) against the implicit baseline. The ensuing parameters were used for representational similarity analysis (RSA) (44, 59).

#### Whole brain searchlight analysis: Node coding

To identify brain regions that encode representations of individual people and places (i.e., the nodes of the schemas), we employed an RSA searchlight analysis (spheres with a radius of 8mm, 4 voxels) across all gray matter voxels. This analysis was based on the RSA toolbox 59 and compared activity patterns across functional runs (54, 96). It identified regions where two simulations of the same person or place yielded more similar activity patterns (same-item similarity) than any two simulations of different people or places (different-item similarity). Specifically, we assessed same-item similarity as the Pearson correlation between the activity pattern of the initial simulation of any given node in the first and its repeated simulation in the second run. Different-item similarity was computed as the average correlation of the initial simulation of a node in the first run with all other nodes of the same category (people or places) in the second run. By constraining the different-item similarity to items of the same category, we ensure that it is not affected by general differences in the neural representation of people versus places. Finally, we determined the magnitude of the node coding as the difference score between same- and different-item similarity (19, 54).

This searchlight analysis yielded a node-coding map for each individual participant. For second level analyses, we *Fisherz*-transformed these maps, normalized them into MNI space using the DARTEL (97) estimated deformation fields, and smoothed them with a Gaussian Kernel of 8 mm radius at full-width-half-maximum. We then masked the smoothed map with the normalized gray matter masks and tested the significance of the node-coding effect using a simple *t*-contrast at each voxel. We used voxel-level inference at *p* < 0.05 (family-wise-error-corrected) and regarded only clusters that comprised at least 30 contiguous voxels.

#### ROI-based analyses: Examining the edges

The second RSA examined whether regions that code for the nodes of the schema also code for the predicted relationships between the nodes (i.e., their edges). This analysis thus examined data from regions-of-interest based on the thresholded node-coding map. Note that the two sets of analyses are based on different parts of the neural RSM and on comparisons of model RSMs that are independent from the node-coding model.

For the mPFC, we joined the two rostral and ventral clusters. For the PCC, we took the conjunction of a broad cluster that included this region and an anatomical PCC mask from the Brainnetome atlas (58) (regions 175, 176, 181, 182). For the hippocampal ROI, we merged its rostral and caudal parts of the same atlas (regions 215 – 218). Voxels were included if they had at least 50% probability of being part of the mask and gray matter.

As complementary analyses, we also examined the data solely based on an anatomical mask of the ventral mPFC used by ref. 19 (see also 98), a more spatially extended mask including the rostral mPFC (comprising Brainnetome regions 13, 14, 41, 42, 47 – 50, 187, 188), and said anatomical mask of the PCC. All masks were inverse normalized into subject space using the DARTEL estimated deformation fields and constrained using the implicit mask estimated from the first level GLMs.

#### Extraction of the importance weights

We had hypothesized that centrality, experience, and affective value would jointly contribute to the importance of a node and expected that they would share a common latent factor. We thus applied principal component analysis (PCA) to the three features and computed the latent factor that explained the most variance. The PCAs were conducted separately for people and places and were based on values of each variable that had been *z*-scored for each participant. This approach ascertained that neither between-category variance nor between-participant variance would bias the factor solution. We then extracted, across all participants, the respective first principal component for people and places. These principal components were positively correlated not only with centrality and experience but also with affective value, consistent with our proposal that all three contributing features jointly quantify the importance of a given node. We thus refer to these principal components as importance factors.

#### Predicting the structure of the edges

We used the importance values to predict the structure of schematic representations in the mPFC. We had hypothesized that more important nodes should, overall, exhibit greater neural similarity with the other nodes. We thus predicted the similarity for any pair of nodes by the product of their respective importance values. We scaled the vectors to the interval of zero (lowest importance) and one (highest importance) prior to multiplication. We then arranged the combined importance values in square matrices for each category (people, places). In initial analyses, we examined the structure of representations separately for the schemas comprising people and places. Note that all analyses are only based on the lower triangular vector of the representational similarity matrices.

#### Model comparisons using Linear Mixed Models

We set up a series of linear mixed effects models in R (version 3.5.1, www.r-project.org), using LME4 (99), to test which of several alternative predictors accounted best for the structure of representations in the ROIs. These models accounted for the neural similarity data as a function of the full fixed effects of category (people, places) and predictor of interest (i.e., centrality, experience, affective value, or the principal component). We further accounted for between participant variance by including random effects: one random intercept for participant and run as well as random slopes for category and the respective predictor. Hence, all models were of the form:

~~~
Neural similarity ∼ category * predictor + (1 + category + predictor|participant) + (1|participant:run)
~~~

We estimated the models separately for each ROI and subsequently performed model comparisons based on the relative Log Evidence Ratios (*LER*) derived from Akaike’s Information Criterion (60). The best model assumes, by definition, a relative *LER* of zero, and we regard relative *LER* differences greater than two as *decisive* evidence for the better model (61).

We further examined whether the winning models in each ROI are also substantially superior to models based on random Gaussian noise. We thus created null models by randomly sampling 30 values from a standard normal distribution for both people and places. We then rescaled these values to the interval from zero to one. Subsequently, we constructed a noise null model by computing the product of every combination of two values, just as we had done for our predictors of interest. We also created a second null model by sorting the same random noise values in descending order prior to multiplication. This was done to account for the inherent order of the original lists of people and places provided by the participants that tended to start with more familiar and pleasant exemplars. Thus, people and places that were named first always received larger random numbers than those named later.

We then fit linear mixed effect models for these two noise null models, and performed a model comparison with the winning model(s) from the respective ROI. We repeated this estimation process 1,000 times to compute average model performance. Critically, if the winning model(s) in each ROI constitute(s) a good approximation of the structure of neural representations, they should consistently outperform both the random noise and the sorted noise models.

## ACKNOWLEDGEMENTS

This work was supported by a Max Planck Research Group (awarded to RGB). We thank Roxanne Eisenbeis for assistance in data collection, Ruud Berkers for help in setting up preprocessing, Davide Stramaccia for discussing the linear mixed effects models, and Angharad Williams for comments on an earlier draft of this manuscript.

## AUTHOR CONTRIBUTIONS

*Conceptualization:* P.C.P. and R.G.B.; *Methodology:* P.C.P, I.C., and R.G.B.; *Investigation:* P.C.P; *Software:* P.C.P. and I.C.; *Formal Analysis:* P.C.P, I.C., and R.G.B.; *Visualization:* P.C.P; *Writing – Original Draft:* P.C.P. and R.G.B.; *Writing – Review & Editing:* P.C.P, I.C., and R.G.B.; *Funding Acquisition:* R.G.B.; *Supervision:* R.G.B.

