## Supplement for "Value shapes the structure of schematic representations in the medial prefrontal cortex"

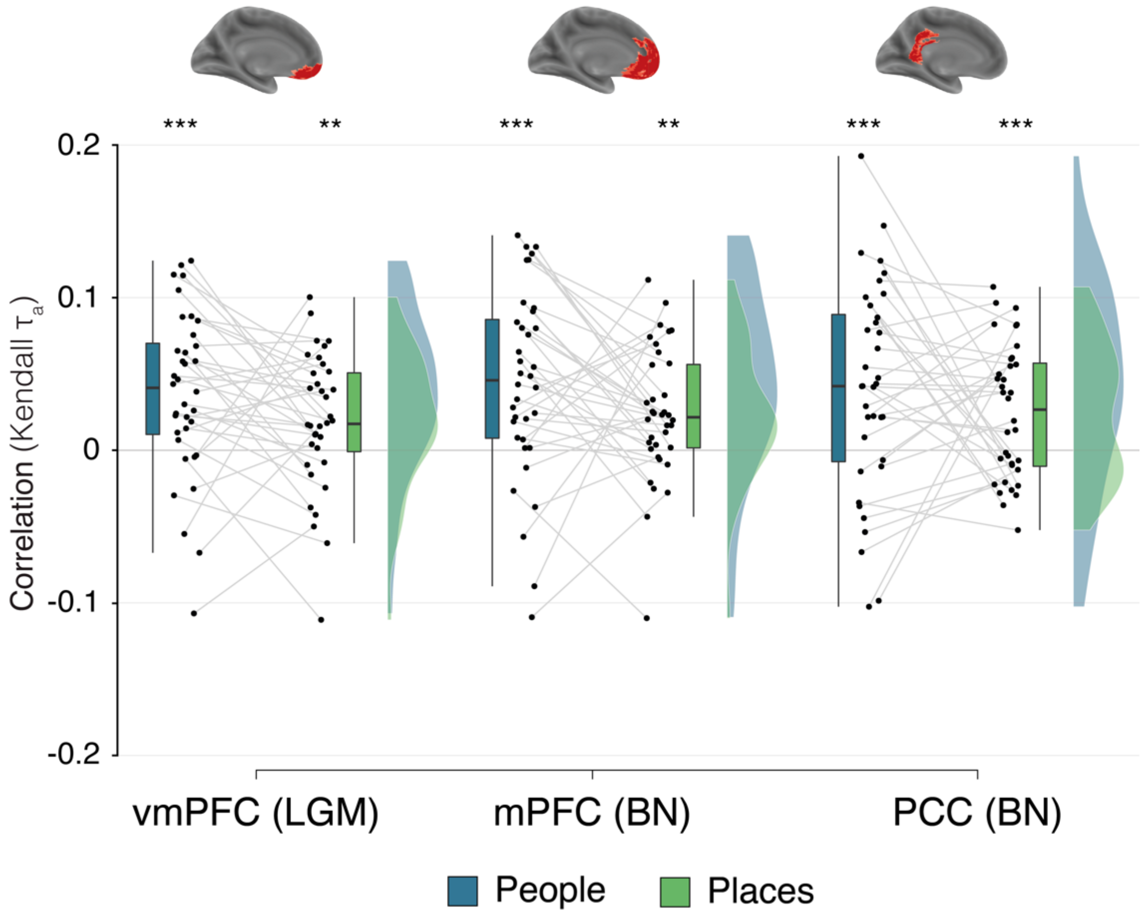


Figure S1. Correlation of the principal component in anatomical ROIs.

Asterisks denote the result of a directed *t*-test on the *Fisher-Z* transformed correlation coefficients (*** - $p < .001$, ** - $p < .01$, vmPFC = ventromedial prefrontal cortex, mPFC = medial prefrontal cortex, PCC = posterior cingulate cortex, LGM = Liu, Grady, Moscovitch (see also Benoit et al., 2019; Liu et al., 2017), BN = Brainnetome (Fan et al., 2016)). Box-plots: center line, median; box limits, first and third quartile; whiskers, 1.5x interquartile range.

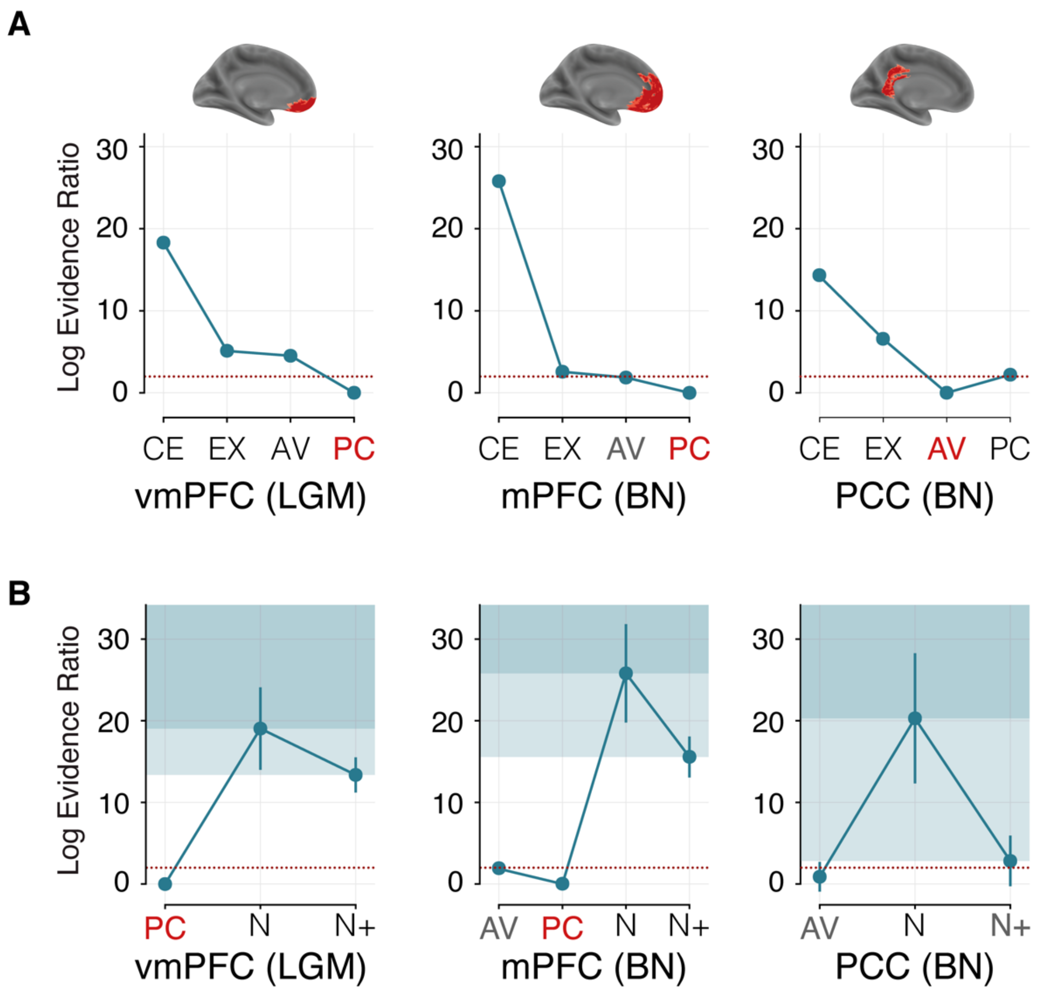


Figure S2. Linear mixed effects models – Model selection in anatomical ROIs.

(A) Model comparison in three additional, anatomical ROIs, (B) Model comparisons of the winning models against random gaussian noise and sorted gaussian noise. Data points depict the mean model performance in comparisons with 1,000 random noise models (N) and sorted noise models (N+), whiskers indicate the standard deviation of the model performance. vmPFC = ventromedial prefrontal cortex, mPFC = medial prefrontal cortex, PCC = posterior cingulate cortex, LGM = Liu, Grady, Moscovitch (see also Benoit et al., 2019; Liu et al., 2017), BN = Brainnetome (Fan et al., 2016), CE = centrality, EX = experience, AV = affective value, PC = principal component, N = random noise, N+ = sorted noise.

Table S1. Searchlight analysis – Node coding

| Region (peak) | approx. BA (peak) | Hemi-sphere | MNI (peak) *x* | *y* | *z* | Voxels | *Z(max)* |
|---|---|---|---|---|---|---|---|
| PCC, precuneus | 7, 23, 31 | L/R | -2 | -58 | 24 | 10,335 | 7.19 |
| | | | -2 | -50 | 28 | | 7.14 |
| | | | -2 | -62 | 42 | | 7.08 |
| Medial PFC | 10, 11 | L | -4 | 56 | 2 | 353 | 6.54 |
| | | | -10 | 68 | 0 | | 5.30 |
| | | | -8 | 44 | -14 | | 5.04 |
| Dorsolateral PFC | 45, 46 | L | -52 | 34 | 10 | 289 | 6.48 |
| | | | -46 | 32 | 18 | | 5.58 |
| | | | -40 | 38 | 18 | | 5.41 |
| Lateral PFC | 6, 44 | L | -48 | 8 | 32 | 297 | 6.38 |
| | | | -42 | 4 | 44 | | 5.71 |
| | | | -52 | 16 | 26 | | 5.07 |
| Dorsal mPFC | 9, 10 | R | 4 | 50 | 20 | 227 | 6.03 |
| | | | 10 | 56 | 28 | | 5.57 |
| | | | 4 | 58 | 18 | | 5.40 |
| Fronto-parietal cortex | 6, 8 | L | -26 | 24 | 46 | 401 | 5.92 |
| | | | -22 | 24 | 56 | | 5.40 |
| | | | -28 | 4 | 58 | | 5.35 |
| Early visual | visual assoc | L | 16 | -90 | 0 | 133 | 5.67 |
| Fronto-parietal cortex | 6 | R | 34 | 8 | 58 | 148 | 5.66 |
| | | | 40 | 2 | 50 | | 5.48 |
| | | | 26 | 12 | 58 | | 5.46 |
| Early visual | visual assoc | L | -12 | -92 | -2 | 53 | 5.55 |
| | | | -6 | -98 | -6 | | 5.13 |
| Lateral PFC | 44, 45, 46 | R | 58 | 22 | 18 | 146 | 5.54 |
| | | | 56 | 24 | 10 | | 5.51 |
| | | | 44 | 38 | 10 | | 5.20 |
| Dorsolateral PFC | 8 | R | 32 | 28 | 40 | 65 | 5.43 |
| | | | 32 | 22 | 48 | | 4.91 |

| Region (peak) | approx. BA (peak) | Hemi-sphere | MNI (peak) *x* | *y* | *z* | Voxels | *Z(max)* |
|---|---|---|---|---|---|---|---|
| Medial PFC | 10 | R | 6 | 54 | -8 | 65 | 5.35 |
| | | | 12 | 50 | -4 | | 4.99 |
| Temporal cortex | 21 | L | -56 | -20 | -10 | 40 | 5.34 |
| Lateral PFC | 9 | R | 36 | 42 | 24 | 50 | 5.21 |
| | | | 42 | 36 | 20 | | 5.15 |
| Temporal cortex | 21, fusiform | R | 64 | -44 | 4 | 137 | 5.17 |
| | | | 56 | -46 | -6 | | 5.17 |
| | | | 58 | -52 | 2 | | 5.13 |
| Dorsal mPFC | 9, 10 | L | -20 | 50 | 28 | 38 | 5.16 |
| | | | -14 | 54 | 32 | | 5.12 |

***Note.*** Thresholded at $p < .05$, voxel FWE corrected and at least 30 contiguous voxels. BA = Brodmann area.

Table S2. Correlation of the principal component in anatomical ROIs

| | | Descriptive statistics | | Shapiro Wilk test | | Significance test | | |
|---|---|---|---|---|---|---|---|---|
| ROI | Category | *Mean corr.* | *SD* | *W* | *p* | *t(35)* | *p* | *d* |
| vmPFC (LGM) | people | 0.037 | 0.054 | 0.97 | 0.47 | 4.16 | < .001 | 0.69 |
| | places | 0.019 | 0.045 | 0.97 | 0.41 | 2.52 | 0.008 | 0.42 |
| mPFC (BN) | people | 0.044 | 0.062 | 0.97 | 0.35 | 4.26 | < .001 | 0.71 |
| | places | 0.023 | 0.043 | 0.96 | 0.21 | 3.25 | 0.001 | 0.54 |
| PCC (BN) | people | 0.040 | 0.069 | 0.99 | 0.96 | 3.47 | < .001 | 0.58 |
| | places | 0.025 | 0.044 | 0.95 | 0.13 | 3.41 | < .001 | 0.57 |

*Note.* ROI = Region of interest, *corr.* = Correlation (Kendall's $\tau_a$), *SD* = standard deviation, *d* = Cohen's *d,* vmPFC = ventromedial prefrontal cortex, mPFC = medial prefrontal cortex, PCC = posterior cingulate cortex, LGM = Liu, Grady, Moscovitch (see also Benoit et al., 2019; Liu et al., 2017), BN = Brainnetome (Fan et al., 2016).

Table S3. Correlations of centrality, experience, and affective value

| | | | Descriptive statistics | | Shapiro Wilk test | | Significance test | | |
|---|---|---|---|---|---|---|---|---|---|
| ROI | Model | Category | Mean corr. | SD | W | *p* | *stat* | *p* | *d* |
| mPFC (SL) | CE | people | 0.007 | 0.065 | 0.96 | 0.24 | 0.67 | 0.252 | 0.11 |
| | | places | -0.001 | 0.040 | 0.97 | 0.44 | -0.18 | 0.571 | 0.03 |
| | EX | people | 0.041 | 0.062 | 0.96 | 0.16 | 3.98 | < .001 | 0.66 |
| | | places | 0.023 | 0.047 | 0.97 | 0.45 | 2.92 | 0.003 | 0.49 |
| | AV | people | 0.036 | 0.061 | 0.96 | 0.30 | 3.49 | 0.001 | 0.58 |
| | | places | 0.022 | 0.051 | 0.97 | 0.38 | 2.61 | 0.007 | 0.43 |
| HPC (BN) | CE | people | -0.013 | 0.035 | 0.98 | 0.59 | -2.23 | 0.984 | 0.37 |
| | | places | -0.001 | 0.036 | 0.94 | *0.04* | 303$^{w}$ | 0.682 | 0.03 |
| | EX | people | 0.005 | 0.041 | 0.97 | 0.31 | 0.68 | 0.25 | 0.11 |
| | | places | 0.006 | 0.037 | 0.97 | 0.39 | 0.92 | 0.182 | 0.15 |
| | AV | people | 0.007 | 0.043 | 0.97 | 0.48 | 1.04 | 0.152 | 0.17 |
| | | places | 0.005 | 0.035 | 0.98 | 0.80 | 0.82 | 0.208 | 0.14 |
| PCC (SL) | CE | people | 0.005 | 0.064 | 0.95 | 0.07 | 0.46 | 0.323 | 0.08 |
| | | places | 0.004 | 0.045 | 0.97 | 0.41 | 0.59 | 0.279 | 0.10 |
| | EX | people | 0.048 | 0.056 | 0.97 | 0.33 | 5.15 | < .001 | 0.86 |
| | | places | 0.024 | 0.049 | 0.97 | 0.54 | 2.93 | 0.003 | 0.49 |
| | AV | people | 0.035 | 0.070 | 0.96 | 0.23 | 3.03 | 0.002 | 0.50 |
| | | places | 0.018 | 0.063 | 0.96 | 0.21 | 1.75 | 0.045 | 0.29 |

| ROI | Model | Category | Descriptive statistics: Mean corr. | Descriptive statistics: SD | Shapiro Wilk test: W | Shapiro Wilk test: *p* | Significance test: *stat* | Significance test: *p* | Significance test: *d* |
|---|---|---|---|---|---|---|---|---|---|
| vmPFC (LGM) | CE | people | 0.003 | 0.051 | 0.98 | 0.75 | 0.40 | 0.347 | 0.07 |
| | | places | -0.004 | 0.039 | 0.86 | < .001 | 373[w] | 0.27 | 0.10 |
| | EX | people | 0.039 | 0.052 | 0.98 | 0.80 | 4.45 | < .001 | 0.74 |
| | | places | 0.017 | 0.041 | 0.98 | 0.86 | 2.50 | 0.009 | 0.42 |
| | AV | people | 0.034 | 0.053 | 0.97 | 0.37 | 3.84 | < .001 | 0.64 |
| | | places | 0.015 | 0.049 | 0.97 | 0.49 | 1.91 | 0.032 | 0.32 |
| mPFC (BN) | CE | people | 0.003 | 0.057 | 0.98 | 0.72 | 0.29 | 0.387 | 0.05 |
| | | places | 0.001 | 0.043 | 0.87 | *<.001* | 388[w] | 0.198 | 0.01 |
| | EX | people | 0.049 | 0.057 | 0.97 | 0.36 | 5.18 | < .001 | 0.86 |
| | | places | 0.021 | 0.045 | 0.98 | 0.60 | 2.81 | 0.004 | 0.47 |
| | AV | people | 0.037 | 0.061 | 0.98 | 0.71 | 3.68 | < .001 | 0.61 |
| | | places | 0.018 | 0.050 | 0.97 | 0.36 | 2.17 | 0.018 | 0.36 |
| PCC (BN) | CE | people | -0.001 | 0.061 | 0.94 | 0.07 | -0.12 | 0.548 | 0.02 |
| | | places | 0.007 | 0.050 | 0.98 | 0.87 | 0.88 | 0.191 | 0.15 |
| | EX | people | 0.043 | 0.058 | 0.98 | 0.84 | 4.40 | < .001 | 0.73 |
| | | places | 0.026 | 0.047 | 0.99 | 0.97 | 3.35 | 0.001 | 0.56 |
| | AV | people | 0.032 | 0.071 | 0.96 | 0.22 | 2.70 | 0.005 | 0.45 |
| | | places | 0.013 | 0.060 | 0.96 | 0.17 | 1.35 | 0.092 | 0.23 |

***Note.*** [w] – statistic *W* of a Wilcox test, used due to a deviation from normality as indicated by the Shapiro-Wilk-Test. All other statistics indicate the *t*-statistic of a simple *t*-test ($df = 35$). mPFC = medial prefrontal cortex, HPC = hippocampus, PCC = posterior cingulate cortex, vmPFC = ventromedial prefrontal cortex, SL = searchlight, LGM = Liu, Grady, Moscovitch (see also Benoit et al., 2019; Liu et al., 2017), BN = Brainnetome (Fan et al., 2016), CE = centrality, EX = experience, AV = affective value, Mean corr. = Mean correlation (Kendall's $\tau_a$), SD = standard deviation, *d* = Cohen's *d*.

Table S4. Linear mixed effects models – Model parameters of the winning models

| Region | Winning model(s) | Effect | *β* | *SE* | $\chi^2$ | *p* | *Sig.* |
|---|---|---|---|---|---|---|---|
| mPFC (SL) | PC | $\text{Category}_{place}$ | -0.026 | 0.008 | 11.61 | < .001 | *** |
| | | PC | 0.048 | 0.012 | 17.12 | < .001 | *** |
| | | $\text{Category}_{place}$:PC | -0.005 | 0.008 | 0.36 | 0.546 | *n.s.* |
| HPC (BN) | CE | $\text{Category}_{place}$ | 0.020 | 0.004 | 31.49 | < .001 | *** |
| | | CE | -0.007 | 0.004 | 0.90 | 0.342 | *n.s.* |
| | | $\text{Category}_{place}$:CE | 0.007 | 0.004 | 3.04 | 0.081 | *n.s.* |
| | AV | $\text{Category}_{place}$ | 0.023 | 0.004 | 31.47 | < .001 | *** |
| | | AV | 0.004 | 0.004 | 1.47 | 0.225 | *n.s.* |
| | | $\text{Category}_{place}$:AV | -0.002 | 0.004 | 0.16 | 0.686 | *n.s.* |
| PCC (SL) | AV | $\text{Category}_{place}$ | 0.014 | 0.009 | 1.51 | 0.219 | *n.s.* |
| | | AV | 0.026 | 0.007 | 10.71 | 0.001 | ** |
| | | $\text{Category}_{place}$:AV | -0.010 | 0.005 | 3.92 | 0.048 | * |
| vmPFC (LGM) | PC | $\text{Category}_{place}$ | -0.019 | 0.005 | 19.30 | < .001 | *** |
| | | PC | 0.027 | 0.007 | 11.75 | < .001 | *** |
| | | $\text{Category}_{place}$:PC | -0.009 | 0.006 | 2.28 | 0.131 | *n.s.* |
| mPFC (BN) | PC | $\text{Category}_{place}$ | -0.011 | 0.006 | 5.29 | 0.021 | * |
| | | PC | 0.037 | 0.007 | 23.62 | < .001 | *** |
| | | $\text{Category}_{place}$:PC | -0.008 | 0.006 | 1.56 | 0.211 | *n.s.* |
| | AV | $\text{Category}_{place}$ | -0.010 | 0.006 | 4.32 | 0.038 | * |
| | | AV | 0.025 | 0.006 | 14.35 | < .001 | *** |
| | | $\text{Category}_{place}$:AV | -0.005 | 0.005 | 0.97 | 0.326 | *n.s.* |
| PCC (BN) | AV | $\text{Category}_{place}$ | 0.057 | 0.008 | 52.72 | < .001 | *** |
| | | AV | 0.021 | 0.007 | 6.87 | 0.009 | ** |
| | | $\text{Category}_{place}$:AV | -0.010 | 0.005 | 4.84 | 0.028 | * |

*Note.* mPFC = medial prefrontal cortex, HPC = Hippocampus, PCC = posterior cingulate cortex, vmPFC = ventromedial prefrontal cortex, SL = searchlight, BN = Brainnetome (Fan et al., 2016), LGM = Liu, Grady, Moscovitch (see also Benoit et al., 2019; Liu et al., 2017), CE = centrality, EX = experience, AV = affective value, PC = principal component, SE = standard error, Sig. = significance (***– $p < .001$, ** – $p < .01$, * – $p < .05$).

Table S5. Searchlight analysis – Principal component

| Region (peak) | approx. BA (peak) | Hemi-sphere | MNI (peak) x | y | z | Voxel | *Z(max)* |
|---|---|---|---|---|---|---|---|
| Medial PFC | 6, 8, 9 | L/R | -4 | 32 | 50 | 9,375 | 5.91 |
| | | | 18 | 28 | 58 | | 5.24 |
| | | | 10 | 58 | 30 | | 5.16 |
| Dorsolateral PFC | 6, 8, 45 | L | -52 | 24 | 8 | 1,189 | 4.79 |
| | | | -38 | 14 | 48 | | 4.72 |
| | | | -44 | 20 | 40 | | 4.24 |
| Lateral parietal cortex | 19, 39 | L | -38 | -60 | 20 | 1,087 | 4.70 |
| | | | -50 | -68 | 38 | | 4.34 |
| | | | -54 | -64 | 30 | | 4.30 |
| Lateral PFC | 44 | R | 54 | 18 | 14 | 420 | 4.68 |
| | | | 58 | 14 | 8 | | 4.22 |
| | | | 50 | 10 | -2 | | 3.58 |
| Cerebellum | - | L/R | 38 | -60 | -40 | 2,005 | 4.44 |
| | | | 34 | -64 | -32 | | 4.32 |
| | | | 34 | -62 | -48 | | 4.26 |
| posterior cingulate cortex | 23 | L/R | 2 | -44 | 36 | 1,126 | 4.40 |
| | | | 4 | -46 | 28 | | 4.33 |
| | | | -4 | -40 | 26 | | 4.03 |
| Lateral PFC | 45, 47 | R | 38 | 34 | -10 | 451 | 4.28 |
| | | | 46 | 30 | -10 | | 3.99 |
| | | | 52 | 30 | 4 | | 3.43 |
| Lateral parietal cortex | 39 | R | 58 | -56 | 28 | 725 | 4.26 |
| | | | 48 | -46 | 28 | | 4.04 |
| | | | 50 | -54 | 34 | | 3.95 |
| Lateral temporal cortex | 22, 41 | R | 60 | -20 | 4 | 246 | 4.12 |
| | | | 60 | -22 | -4 | | 3.75 |
| Lateral temporal cortex | 21 | L | -62 | -10 | -24 | 59 | 3.99 |
| Anterior temporal cortex | 20, 21 | R | 56 | -6 | -36 | 224 | 3.94 |
| | | | 52 | -2 | -28 | | 3.60 |

| Region (peak) | approx. BA (peak) | Hemi-sphere | MNI (peak) | | | Voxel | *Z(max)* |
|---|---|---|---|---|---|---|---|
| | | | *x* | *y* | *z* | | |
| Temporal cortex | 21, 22 | L | -54 | -24 | -10 | 214 | 3.89 |
| | | | -58 | -26 | 0 | | 3.51 |
| | | | -54 | -32 | -4 | | 3.44 |
| Anterior temporal cortex | 38 | L | -50 | -2 | -32 | 58 | 3.84 |
| Lateral parietal | 40 | L | -46 | -32 | 14 | 76 | 3.79 |
| | | | -48 | -22 | 12 | | 3.36 |
| Anterior insula | 47 | L | -32 | 28 | -8 | 134 | 3.77 |
| | | | -38 | 22 | -12 | | 3.71 |
| | | | -30 | 20 | -28 | | 3.62 |
| Thalamus | - | L | -4 | -34 | 6 | 44 | 3.63 |
| Lateral PFC | 47 | L | -48 | 42 | -2 | 37 | 3.52 |
| Cerebellum | - | L | -42 | -66 | -54 | 51 | 3.49 |
| Ventromedial PFC | 11 | R | 8 | 24 | -16 | 32 | 3.43 |

***Note.*** For exploratory purposes, thresholded at $p < 0.001$, uncorrected, and at least 30 contiguous voxels. BA = Brodmann area.
